# Modulation of B cell receptor activation by antibody competition

**DOI:** 10.1101/2024.11.18.624200

**Authors:** Yuanyuan He, Zijian Guo, Michael D. Vahey

## Abstract

During repeated virus exposure, pre-existing antibodies can mask viral epitopes by competing with B cell receptors for antigen. Although this phenomenon has the potential to steer B cell responses away from conserved epitopes, the factors that influence epitope masking by competing antibodies remain unclear. Using engineered, influenza-reactive B cells, we investigate how antibodies influence the accessibility of epitopes on the viral surface. We find that membrane-proximal epitopes on influenza hemagglutinin are fundamentally at a disadvantage for B cell recognition because they can be blocked by both directly and indirectly competing antibodies. While many influenza-specific antibodies can inhibit B cell activation, the potency of masking depends on proximity of the targeted epitopes as well as antibody affinity, kinetics, and valency. Although most antibodies are inhibitory, we identify one that can enhance accessibility of hidden viral epitopes. Together, these findings establish rules for epitope masking that could help advance immunogen design.

## INTRODUCTION

Adaptive immunity protects against viral infections by recognizing epitopes on viral surface proteins and eliciting antibodies that bind to them^1^. During the initial exposure to a virus, naïve B cells become activated through the binding of antigen-specific B cell receptors (BCRs) and differentiate into either short-lived plasmablasts or germinal center B cells, the latter of which undergo further clonal expansion and affinity maturation^2–4^. This process results in the generation of both long-lived antibody-secreting plasma cells that produce high affinity and sometimes protective antibodies, as well as memory B cells, which may recognize conserved viral epitopes via their BCR upon subsequent exposures. Recalled memory B cells have the potential to generate a rapid and more potent immune response than naïve B cells through differentiation into plasma cells that generate high-affinity antibodies. However, pre-existing antibodies from prior antigen exposure can impede potent recall responses by blocking conserved epitopes through competitive binding. This phenomenon, commonly referred to as epitope masking, can compromise efforts to generate a protective immune response to curb viral infection, as masked epitopes are not efficiently recognized by memory B cells^5–10^.

Evidence of epitope masking has been observed for viral, bacterial, and parasitic infections^5–13^. This phenomenon may be particularly consequential for influenza viruses, where continual viral evolution limits the duration of immunity, and encounters with the virus throughout life are frequent^5–7,10,14,15^. Hemagglutinin (HA), the target of traditional influenza vaccines, is comprised of a poorly-conserved globular head domain which is immunodominant over the more highly-conserved stalk^16,17^. Consequently, antibodies against the head domain often lack cross-reactivity against emerging viral variants while stalk-binding antibodies are more resilient in the presence of antigenic drift. During repeated exposure to influenza A viruses, pre-existing antibodies against the conserved stalk could subdue the activation of memory B cells that recognize overlapping epitopes^18^. In this way, epitope masking has been proposed to contribute to a negative feedback loop that disadvantages recognition of the HA stalk and compromises the duration of immunity^19^.

Although several studies have identified important consequences of epitope masking, factors that influence the competition between pre-existing antibodies and memory B cells during repeated exposure to viral antigens are not well understood. In particular, it remains unclear whether or how sensitivity to epitope masking differs across viral epitopes, and how it is influenced by the affinity, kinetics, and valency of competing antibodies. To begin addressing these questions, we developed an imaging-based method to study the activation of influenza-specific B cells presented with viral antigens. This method allowed us to vary the specificity of antibodies and BCRs and to test how they compete. In addition to observing intrinsic differences in activation levels for B cells targeting epitopes across HA and NA, we found that antibody opsonization of viruses frequently inhibits B cell activation, independent of Fc-mediated signaling. This inhibition can be either direct (i.e., where the antibody binds and BCR bind to the same epitope) or indirect, with membrane-proximal epitopes being most sensitive to masking. While antibody affinity is important, we find that the kinetics of dissociation play a dominant role, with slow dissociation leading to stronger BCR inhibition in a comparison between an affinity/avidity-matched antibody pair. For B cells that recognize the HA trimer interface, we find that activation is sensitive to the stability of the HA trimer and can be either suppressed or enhanced by other antibodies. Surprisingly, we observed that NA-reactive B cells can be inhibited by a subset of anti-HA antibodies, possibly due to steric hindrance from the Fc region. Collectively, these findings provide mechanistic insights into epitope masking that could guide the development of vaccines that are able to overcome constraints from pre-existing immunity.

## RESULTS

### An imaging approach to study activation of engineered influenza-specific B cells

To dissect the rules governing competition between soluble antibodies and membrane-anchored B cell receptors, we used CRISPR/Cas9 to knock out the endogenous IgM BCR from Ramos B cells and transduced them via lentivirus with a single-chain BCR^20^ derived from selected HA-or NA-reactive antibodies (Figure 1A). This approach enables us to precisely control the epitope that each B cell line recognizes, and it defines the BCR affinity towards its target. Comparing these engineered monoclonal antibody-derived (‘emAb’) B cells expressing IgM-isotype BCRs to wildtype Ramos B cells, we observed similar BCR expression levels (Figure S1A). For all remaining experiments, we used emAb cell lines expressing IgG-isotype BCRs to more closely mimic memory B cells that have undergone isotype switching and multiple arounds of affinity maturation after exposure to influenza viral proteins.

**Figure 1.**
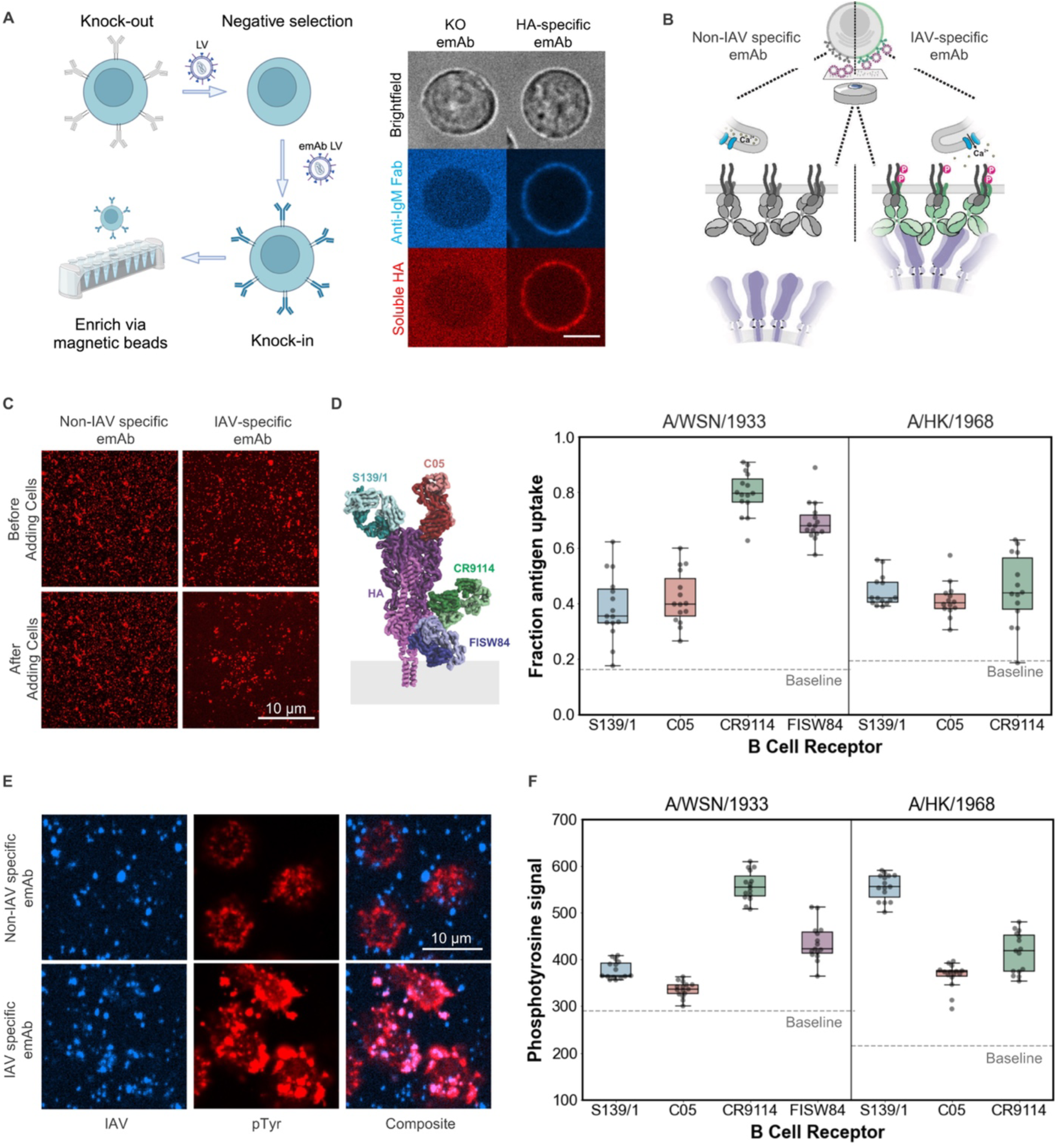
Engineered influenza-specific B cells are activated by surface-bound influenza virus particles. (A) Approach for engineering influenza-specific emAb cells by knocking out the endogenous IgM B cell receptor from Ramos B cells and replacing it with an engineered BCR derived from monoclonal antibodies. Images to the right show knockout Ramos B cells (left column) and CR9114 emAb cells (right column). Scale bar = 5μm. (B) Schematic illustrating the antigen uptake assay. Influenza viruses are reversibly tethered to a glass coverslip and presented to influenza-specific emAb cells. Assay readouts include antigen extraction, phosphotyrosine staining, and calcium imaging. (C) Representative images of A/WSN/1933 virus particles, before and after exposure to IAV-specific emAb cells or non-specific emAb cells. (D) Left: Model of HA (PDB ID 6HJQ) aligned with selected Fabs: S139\1 (4GMS), C05 (4FQR), CR9114 (4FQY), and FISW84 (6HJP). Right: Quantification of antigen extraction by different emAb B cells against A/WSN/1933 virus (left plot) or A/Hong Kong/1968 virus (right plot). (E) Representative images of phosphotyroine localization in IAV-specific or non-specific emAb cells presented with A/WSN/1933 virus particles. Contrast is exaggerated to show colocalization between the two channels. (F) Quantification of phosphotyrosine signal for various emAb cells against A/WSN/1933 (left) or A/Hong Kong/1968 (right) virus particles. Individual data points in panels D and F represent values for separate fields of view. Data are combined from three biological replicates containing five fields of view each. *P*-values are determined by independent t test using the median values for biological replicates.

For emAb cells targeting different influenza epitopes, we used fluorescence microscopy to measure activation against surface-bound influenza A virus particles. For this assay, viral particles are reversibly bound to a glass-bottom plate via *Erythrina cristagalli* lectin (ECL) before introducing emAb cells with defined specificity. Fluorescence imaging allows us to measure particle extraction from the coverslip, as well as calcium influx and BCR phosphorylation (Figure 1B). Optimizing the surface density of ECL allows us to differentiate between IAV-specific emAb cells (with a BCR derived from C05^21^), which robustly extract antigen from the coverslip, and non-specific emAb cells (with a BCR derived from a GFP-reactive antibody^22^, ‘N86’) which do not (Figure 1C, Figure S1B).

### Activation of engineered B cells is antigen specific and sensitive to binding affinity

With this experimental setup, we proceeded to compare antigen extraction and BCR phosphorylation between emAb cells that recognize different HA epitopes. Across four emAb cells with distinct specificities, we observed differences in the extraction of A/WSN/1933 virus particles that does not correlate with their modest (<1.5-fold; Figure S1C) differences in BCR expression (Figure 1D). Stalk-and anchor-specific CR9114^23^-and FISW84^25^-emAb cells show more efficient antigen extraction compared to head-specific S139/1^24^-and C05^21^-emAb cells (Figure 1D, left plot). These trends are consistent with phosphotyrosine levels quantified via immunofluorescence at sites where BCRs colocalize with influenza virus particles (Figure 1E & F, left plot). When the same emAb cells are presented with HAs from A/Hong Kong/1968 for which they have higher (S139/1), similar (C05), or lower (CR9114) affinity compared to A/WSN/1933^21,23,24,26^, we find that both antigen extraction and pTyr levels generally follow BCR affinity (Figure 1D & F).

Phosphorylation of BCR-associated ITAMs depends on the segregation of the phosphatase CD45 from the B cell synapse^27,28^. Signaling via T cell receptors and some Fc receptors depends on the distance between the immune cell membrane and the antigenic surface^29–31^. We reasoned that the membrane-to-membrane distance would differ for the BCRs tested here, from ∼30 nm for C05 down to ∼20 nm for CR9114 (Figure 2A). To determine if this results in differences in CD45 exclusion between the BCRs, we imaged antigen accumulation and CD45 exclusion at interfaces between emAb cells and supported bilayers decorated with purified HA from A/Hong Kong/1968 (Figure 2B). While non-HA reactive N86-emAb cells form small, irregularly shaped clusters of HA (likely due to HA binding to sialylated proteins on the B cell surface^32^), S139/1-, C05-and CR9114-emAb lead to more cell spreading and accumulation of antigen into a larger synapse (Figure 2C). Despite the predicted structural differences between the BCR:HA complexes, we find that CD45 exclusion is similar in all cases (Figure 2C, bottom), consistent with the tendency for all three BCRs to become phosphorylated upon engagement with antigen.

**Figure 2.**
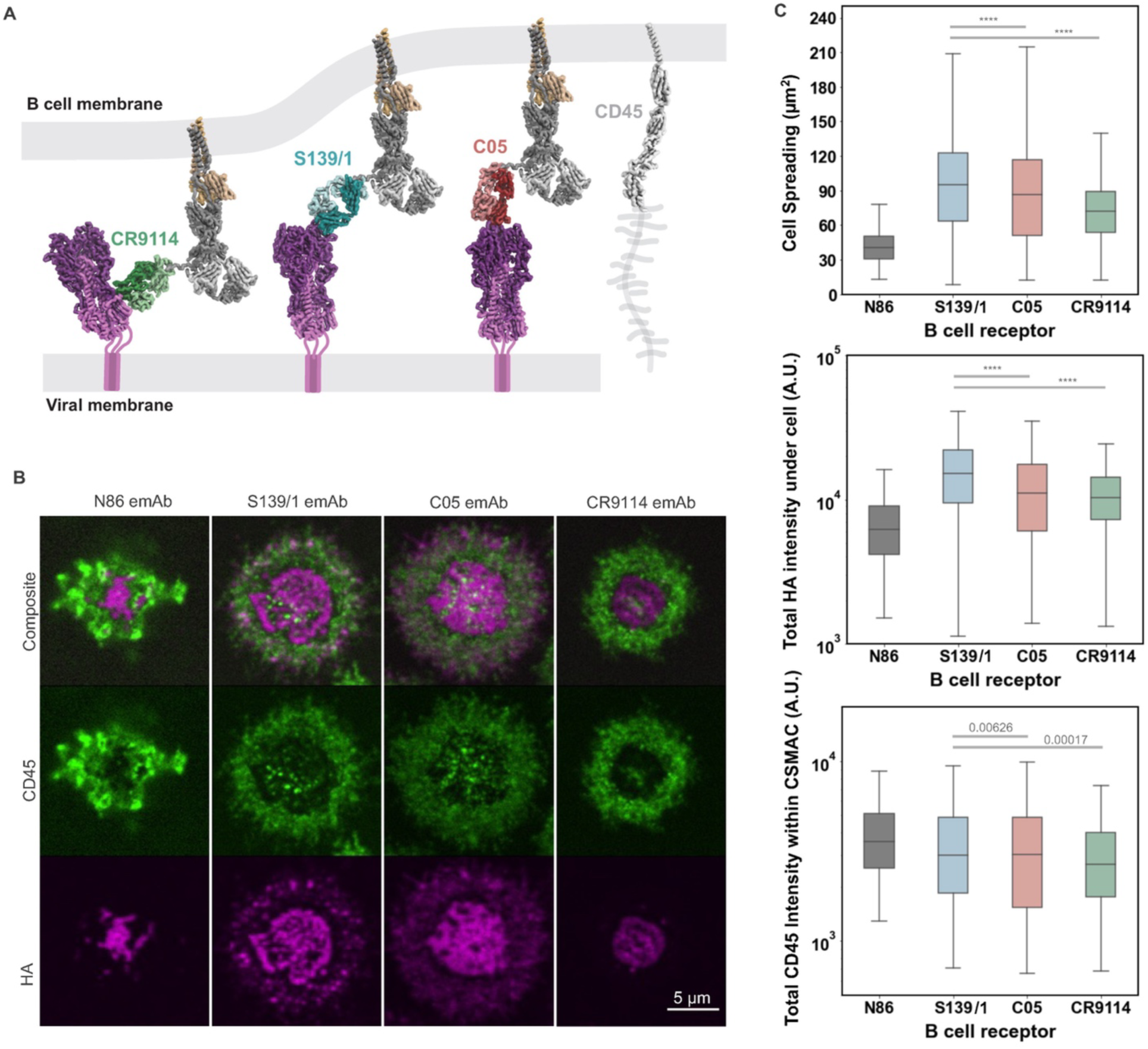
Engineered B cells accumulate HA and exclude CD45 on supported lipid bilayers. (A) Model of BCR engagement with HA. CD45 with a hypothetical depiction of its mucin-like domain is included for comparison. (B) Representative images of emAb cells forming synapses with HA presented on a supported lipid bilayer. (C) Quantification of emAb cell synapses as shown in *B*. Cell spreading (top), HA accumulation within the total footprint of each cell (middle), and CD45 signals within the central supramolecular activation cluster (cSMAC; bottom) are quantified. Individual data points represent measurements performed on individual B cells. Data are combined from two biological replicates containing at least 100 cells per replicate. *P*-values are determined by Kolmogorov-Smirnov test.

### emAb B cells are inhibited by direct competition with antibodies, independent of Fc-mediated effector functions

We next sought to understand how antibodies compete with BCRs during antigen encounter. We first compared engagement of CR9114-emAb cells with A/WSN/1933 virus particles in the presence or absence of directly-competing CR9114 IgG (Supplementary movies S1 and S2). CR9114 IgG almost completely abolished antigen uptake by CR9114-emAb cells and significantly reduced pTyr levels (Figure 3A). We reasoned that this inhibition could arise from epitope masking, as well as from inhibitory Fc interactions, *e.g.*, via FcγRIIb^33^. Using a version of CR9114 that cannot bind to Fc receptors (‘LALAPG’^34^), we observed a similar reduction in pTyr levels as with the wildtype antibody, suggesting that epitope masking alone is sufficient to block B cell activation (Figure 3A). Consistent with this observation, we found that < 0.1% of both wildtype and emAb cells expressed the inhibitory Fc_γ_RIIb on the cell surface (Figure S1D). These findings confirm that epitope masking, and not Fc-dependent signaling, is the main cause of B cell inhibition in our experiments. Epitope masking is also reflected in calcium influx measurements, where CR9114-emAb cells exhibit reduced calcium flux from A/California/04/2009 viruses preincubated with CR9114 IgG (Figure S1E).

**Figure 3.**
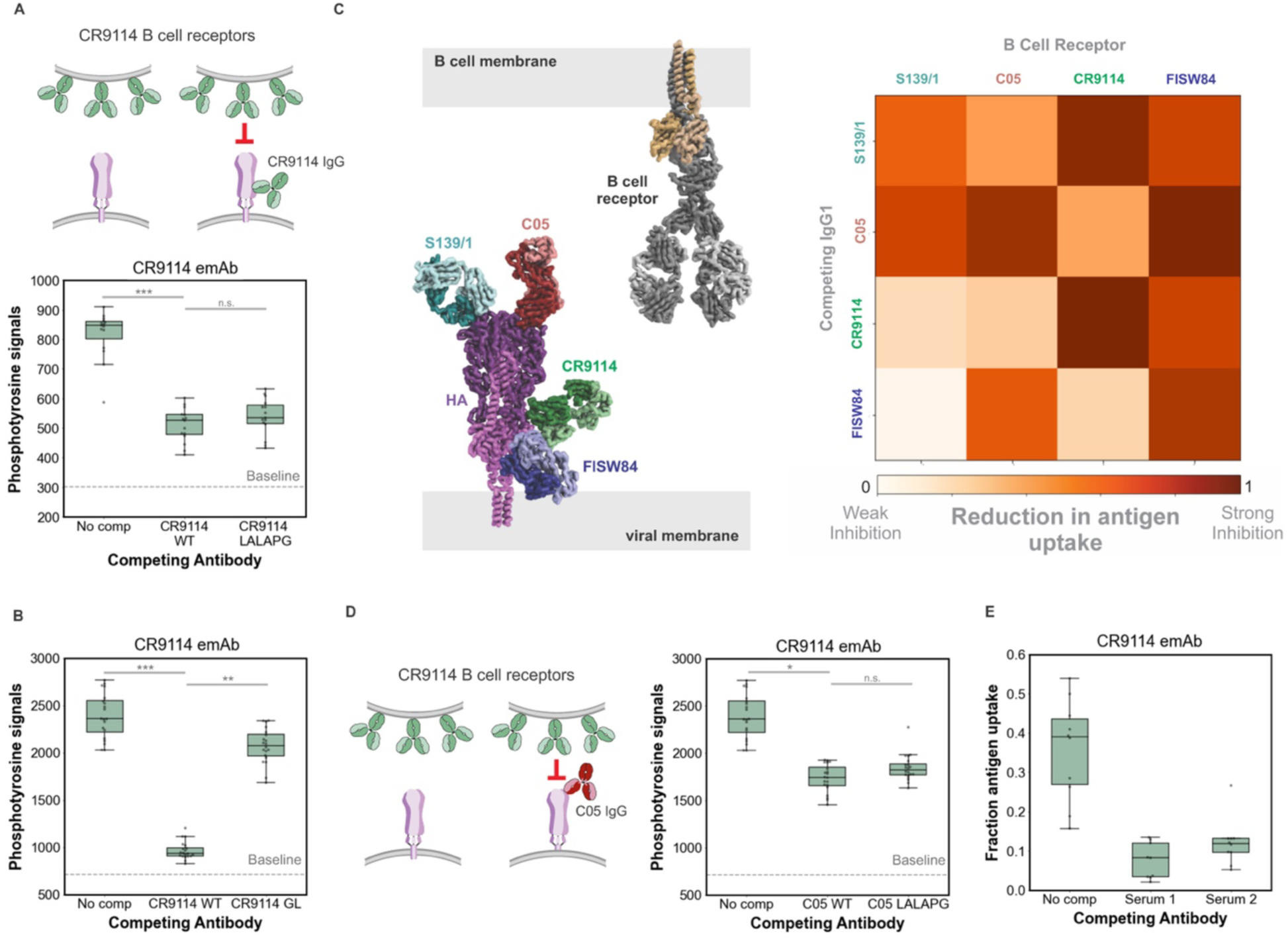
Membrane-proximal epitopes on hemagglutinin are subject to both direct and indirect antibody competition. (A) Competition between CR9114 emAb cells and CR9114 IgG (left cartoon). Plot to the right shows quantification of phosphotyrosine signal from CR9114 emAb cells against A/WSN/1933 viruses with no IgG, 60nM CR9114 IgG, or 60nM CR9114 LALAPG IgG. (B) Quantification of phosphotyrosine signal from CR9114 emAb cells against A/WSN/1933 virus incubated with no IgG, 60nM CR9114 IgG, or 60nM CR9114 IgG reverted to its germline sequence. (C) Direct and indirect competition between HA-specific antibodies and BCRs. Schematic to the left shows a model of the competing antibodies. Plot to the right shows inhibition of antigen uptake for each antibody-BCR pair against A/WSN/1933 virus particles (for C05, CR9114, and FISW84 emAb cells) or A/HK/1963 (for S139 emAb cells). All antibodies are tested at 60nM. (D) Indirect competition between C05 IgG and CR9114 emAb cells. A/WSN/1933 viruses are pre-incubated with no IgG, C05 IgG, or C05 LALAPG IgG at 60nM. (E) Quantification of phosphotyrosine for CR9114 emAb cells against A/California/04/2009 viruses in the presence or absence of purified total IgG from two convalescent sera adjusted to 3.5µM. Data in panels A, B, C, and D are combined from three biological replicates containing five fields of view each. Data in E are combined from two biological replicates containing five fields of view each. *P*-values are determined by independent t test using the median values for biological replicates.

BCRs that have undergone affinity maturation may compete with pre-existing antibodies that bind to the same epitope with lower affinity. To determine how this may effect epitope masking, we reverted CR9114 IgG (K_d_ ∼ 0.4 nM against A/WSN/1933 HA) to its germline sequence (‘CR9114 GL’, K_d_ ∼ 10 nM against A/WSN/1933 HA)^35^. We found that CR9114 GL IgG at 10nM failed to inhibit phosphorylation of CR9114-emAb BCRs, in contrast to the strong inhibition observed for its affinity-matured counterpart (Figure 3B). Thus, in the case of CR9114, affinity-matured B cell receptors can overcome masking by lower-affinity precursor antibodies.

### Indirect antibody competition disadvantages B cells targeting membrane-proximal epitopes

We next asked whether antibodies targeting non-overlapping epitopes could inhibit BCR engagement and B cell activation. For each antibody-BCR pair tested (Figure 3C), we compared antigen extraction in the presence or absence of high concentrations (60 nM) of the competing antibody. We found that in all cases where the antibody and BCR compete for the same epitope, the antibody significantly reduces B cell antigen extraction (Figure 3C, matrix diagonal; Figure S2 A-D). Surprisingly, the head-binding antibodies S139/1 and C05 not only block B cells targeting their respective epitopes, they also potently reduce antigen extraction by both CR9114-and FISW84-emAb cells, whose epitopes lie ∼10 nm away. This effect is recapitulated with soluble S139/1 and CR9114 IgG (Figure S2E). FISW84-emAb cells are inhibited by all antibodies tested, suggesting that sensitivity to epitope masking increases with proximity to the target membrane. However, there are exceptions to this trend: we observe strong inhibition of C05-emAb cells, but not S139/1 emAb cells, by the anchor-targeting FISW84 IgG. Although the mechanism for this inhibition is unclear, it could result from FISW84-induced tilting of the HA ectodomain disrupting binding of the C05 BCR to the HA apex^25,36,37^. Consistent with our previous results, indirect epitope masking does not require Fc effector functions: C05 LALAPG reduces pTyr levels in CR9114-emAb cells to the same extent as normal C05 IgG (Figure 3D). Lastly, we tested the effects of human polyclonal IgG purified from the serum of two individuals infected with A/California/04/2009. We found that both samples tested inhibited extraction of A/California/04/2009 virus by CR9114-emAb cells at ∼3.5µM total IgG (Figure 3E). Collectively, these results demonstrate that a broad spectrum of antibodies can inhibit BCR activation irrespective of overlap between the targeted epitopes, and that BCRs targeting membrane distal epitopes are at a particular disadvantage.

In vivo, B cells may encounter viral antigen in diverse forms, including as particulates or expressed on the surface of antigen presenting cells^38,39^. These formats will differ in both the density and mobility of viral antigen, with potential implications for epitope masking. To investigate if epitope masking occurs when viral antigens can freely diffuse on a fluid membrane, we measured B cell engagement with HA presented on supported lipid bilayers in the presence or absence of competing antibodies (Figure S3A, Supplemental Movie S3). Upon adding either CR9114 or S139/1 IgG, HA from A/Hong Kong/1968 on supported bilayers spontaneously formed diffraction-limited aggregates, likely due to crosslinking by the antibody^40,41^. Subsequent addition of CR9114-emAb cells led to the accumulation of HA aggregates into larger clusters underneath the B cells (Figure S3B). The amount of HA captured as well as CD45 included within the immunological synapse of these cells is smaller in the presence of directly-competing CR9114 IgG, suggesting that epitope masking inhibits B cell engagement with both mobile and particulate antigens. Interestingly, we did not observe inhibition of CR9114 BCRs by indirectly-competing S139/1 IgG on supported bilayers as we had observed for virus particles, suggesting that sensitivity to epitope masking may depend on the manner in which antigen is presented.

### BCR access to the HA trimer interface depends on HA proteolytic activation and can be enhanced or suppressed by antibodies

Conserved epitopes in the HA trimer interface are attractive vaccine targets but may be limited by their reduced accessibility^42–46^. The stability of the HA trimer differs between strains and is increased upon proteolytic activation of HA0 into HA1 and HA2^42,46^. To investigate how these factors affect activation of BCRs targeting the trimer interface, we established emAb cells expressing a BCR derived from an antibody targeting the HA trimer interface (FluA-20^42,46^) and compared phosphotyrosine levels following exposure of these cells to A/WSN/1933 and A/California/04/2009 viruses in both cleaved (HA1/2) and uncleaved (HA0) forms. Despite the conservation of the FluA-20 epitope between these strains, we found stronger activation of FluA-20-emAb cells by A/California/04/2009 compared to A/WSN/1933 HA (Figure 4A), consistent with lower trimer stability in the pandemic H1N1 strain^47^. Moreover, for both strains, HA proteolytic activation with trypsin reduced FluA-20-emAb activation (Figure 4A).

**Figure 4.**
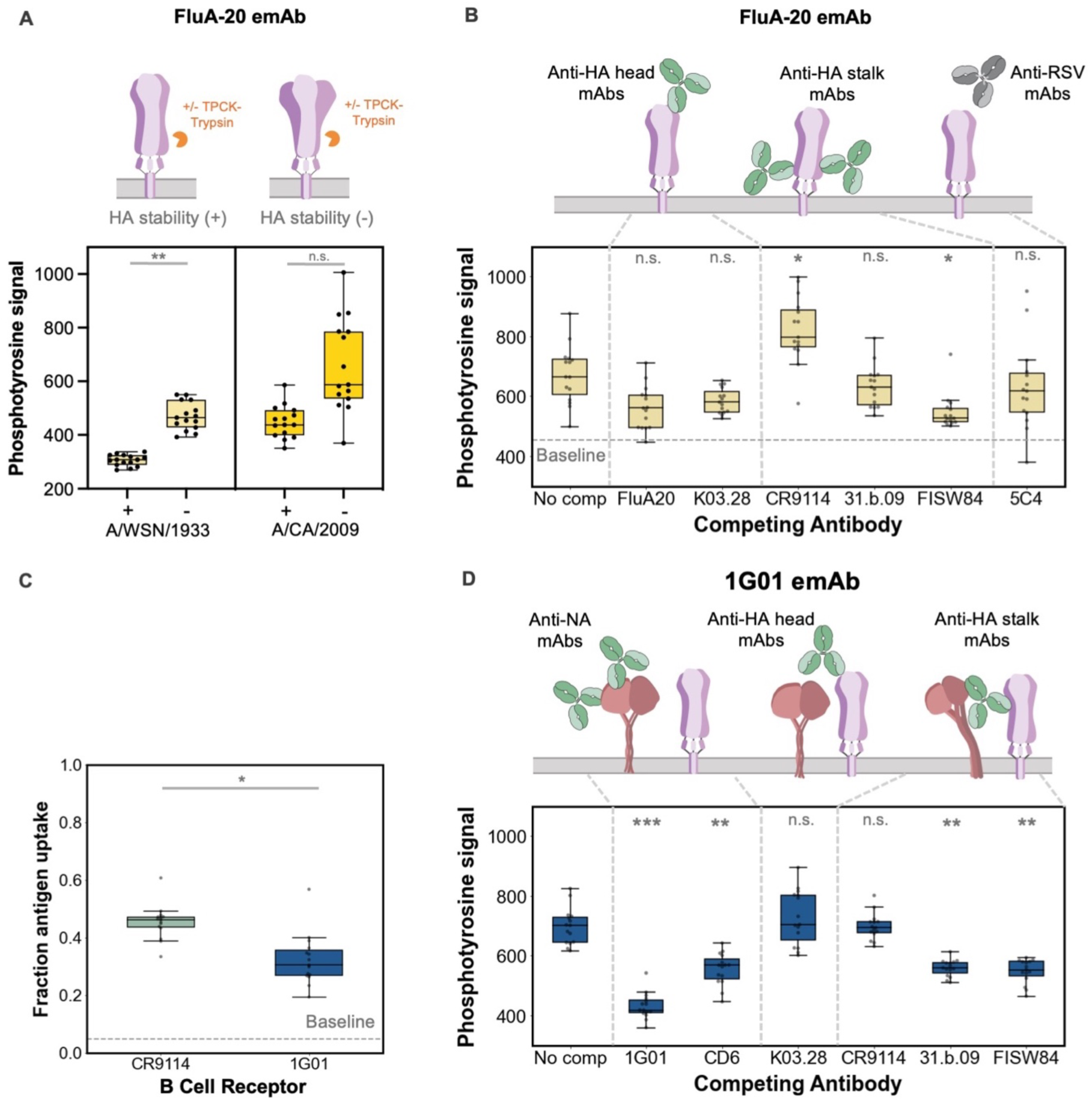
Epitope masking modulates BCR access to the HA trimer interface and NA active site. (A) Quantification of phosphotyrosine signal from FluA-20 emAb cells presented with A/WSN/1933 and A/California/2009 viruses +/-cleavage with TPCK-trypsin. (B) Quantification of phosphotyrosine signal from FluA20 emAb cells against A/California/04/2009 viruses in the presence or absence of the indicated competing IgG (60nM in each case). (C) Comparison of antigen uptake (A/California/04/2009 viruses) by CR9114-and 1G01-emAb cells. (D) Quantification of phosphotyrosine signal from 1G01-emAb RAMOS against A/California/2009 viruses incubated with various mAbs at 60nM. Data in all panels are combined from three biological replicates containing five fields of view each. *P*-values are determined by independent t test using the median values for biological replicates. Statistical comparisons in panels B and D are against the condition without competing IgG.

We next examined how antibody competition influences activation of FluA-20-emAb cells. FluA-20-emAb phosphorylation was reduced by FluA-20 IgG as well as by the head-specific antibody K03.28^48^ and the anchor antibody FISW84, neither of which overlap with the FluA-20 epitope (Figure 4B). Surprisingly, the stalk-specific antibody CR9114 significantly increased the activation of FluA-20-emAb cells, suggesting that it may lead to opening of the trimer. In contrast, 31.b.09^49^, which binds to an epitope in the HA stalk that overlaps with that of CR9114, had little effect on FluA20-emAb activation (Figure 4B). These trends are recapitulated in experiments using competing soluble antibodies, where both CR9114 IgG and Fab led to increased binding by FluA-20 IgG (Figure S3D). Collectively, these results indicate that recognition of the HA trimer interface is highly dependent on trimer stability, the activation status of HA, and can be either suppressed or enhanced in the presence of other antibodies.

### NA-reactive B cells can be inhibited by both anti-HA and anti-NA antibodies

In addition to HA, Neuraminidase (NA) has emerged as an important target of antibody responses to influenza infection^16,18^. To investigate competition between soluble antibodies and NA-reactive emAb cells, we engineered a BCR derived from 1G01, a broadly neutralizing antibody that binds to the NA active site^51^. Despite lower abundance of NA in virus particles relative to HA^52^, 1G01 B cells were still able to extract A/California/04/2009 virions from coverslips, albeit to a lesser extent than CR9114-emAb cells in a side-by-side comparison (Figure 4C). Surprisingly, in antibody competition experiments we found that 1G01 BCRs could be blocked not only by NA-specific 1G01 and CD6^53^, but also by a subset of HA-specific antibodies, including 31.b.09 and FISW84 (Figure 4D). The Fc of antibodies that bind to the HA stalk have previously been reported to inhibit NA enzymatic activity^54^; our results suggest that a similar phenomenon may also influence activation of B cells that bind in or around the NA active site.

### Antibody kinetics and valency regulate the masking potential of antibodies

Antibodies with similar apparent affinities can differ dramatically in underlying binding kinetics. To investigate how this may influence epitope masking, we compared two different antibodies – CR9114 and S139/1 – against HAs where they have similar apparent affinities but widely varying kinetics. While CR9114 IgG binds to A/WSN/1933 HA with slower association and dissociation kinetics, S139/1 IgG binds to A/Hong Kong/1968 HA with high avidity but modest monovalent affinity, leading to more rapid exchange of individual Fabs^23,24^. Both antibodies, however, achieve an apparent *K_d_* of ∼0.4nM. In competition with B cells, we found that CR9114 IgG significantly reduced the activation of CR9114-emAb cells at sub-nanomolar concentration, close to the apparent *K_d_* (Figure 5A, left plot). In contrast, S139/1 IgG required close to 60nM to inhibit S139/1-emAb cells, the maximum concentration tested (Figure 5A, right plot). Thus, despite having similar apparent affinities, antibodies with faster association and dissociation kinetics exhibit a dramatic reduction in masking potency, likely due to faster displacement of bound antibodies with competing BCRs.

**Figure 5.**
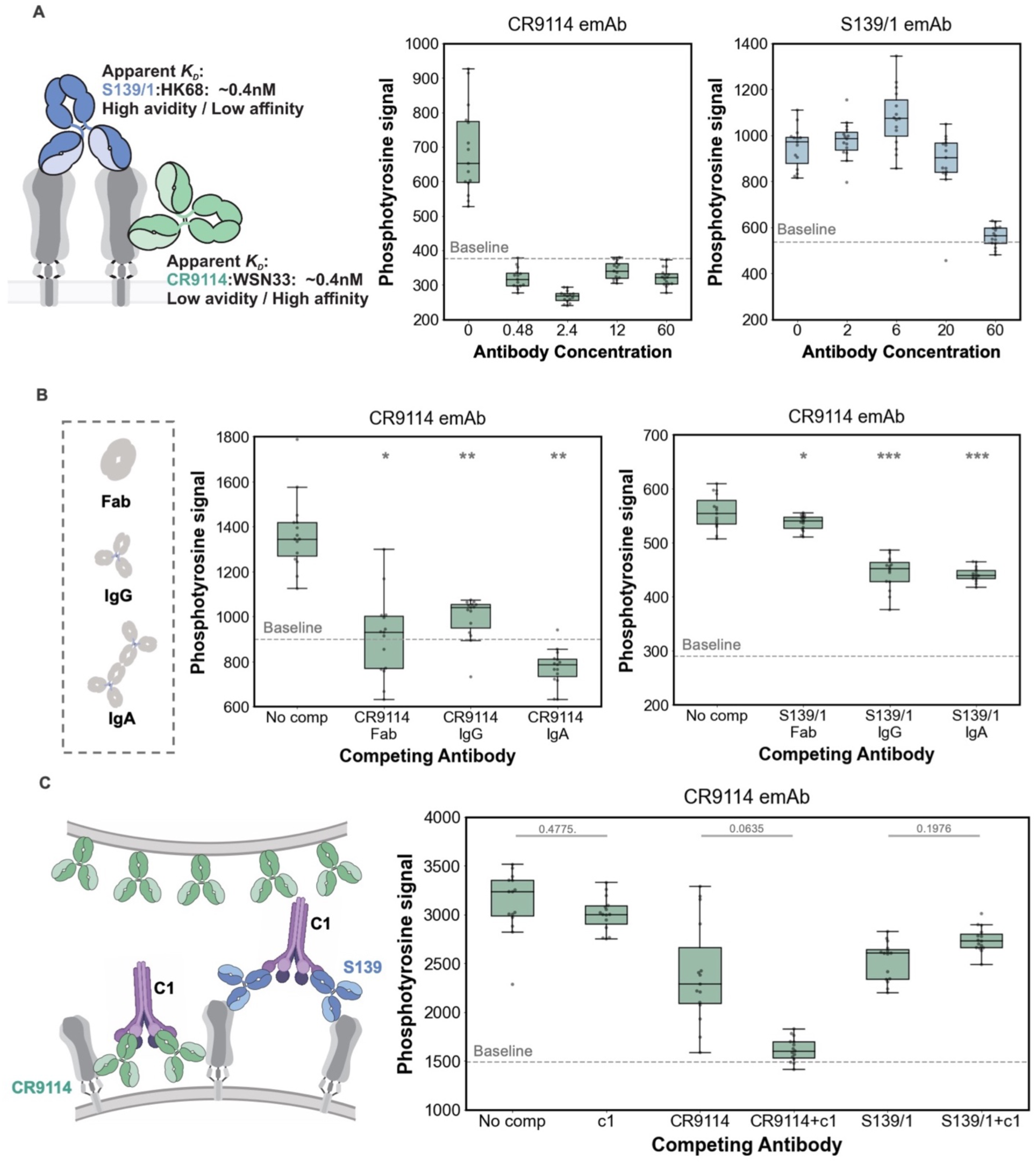
Kinetics, valency, and the binding of complement proteins modulate antibody masking potency. (A) Comparison of epitope masking potency for antibodies with matching apparent affinities but distinct binding kinetics. Plots to the right show quantification of phosphotyrosine signal from CR9114-and S139/1-emAb cells presented with A/WSN/1933 (CR9114) or A/Hong Kong/1968 (S139/1) and increasing concentrations of competing IgG. (B) Left: Quantification of phosphotyrosine signal from CR9114-emAb cells presented with A/WSN/1933 virus particles pre-incubated with CR9114 Fab (120nM), CR9114 IgG (60nM), or CR9114 dIgA (30nM). Right: Quantification of phosphotyrosine signal from CR9114-emAb cells presented with A/WSN/1933 virus particles pre-incubated with S139/1 Fab (120nM), S139/1 IgG (60nM), or S139/1 dIgA (30nM). (C) Quantification of phosphotyrosine signal from CR9114-emAb cells presented with A/WSN/1933 virus particles +/-C1 only, CR9114 IgG +/-C1, or S139/1 +/-C1. For each experiment, IgGs are used at 10nM and C1 is used at 50µg/ml. Data in all panels are combined from three biological replicates containing five fields of view each. *P*-values are determined by independent t test using the median values for biological replicates. Statistical comparisons in panel B are to the condition without competing IgG.

Our observation that head-specific IgGs can block stalk-specific BCRs suggests that steric obstruction is sufficient to achieve epitope masking. To test how increasing antibody size and valency affects epitope masking, we prepared head-specific (S139/1) and stalk-specific (CR9114) antibodies in Fab, IgG and dimeric IgA (dIgA) formats for competition against CR9114-emAb cells at equimolar Fab concentrations (Figure 5B). While S139/1 Fab did not inhibit CR9114-emAb activation, bivalent S139/1 IgG and quadrivalent S139/1 dIgA did, showing similar potency. For stalk-specific CR9114, we found that all formats were similarly potent at blocking CR9114-emAb phosphorylation regardless of size and valency. When CR9114 IgG is combined with complement component 1 (C1), inhibition of CR9114 emAb cells is increased further, resembling the effect of C1 on the neutralizing potency of stalk-binding antibodies (Figure 5C)^55^. In contrast, combining C1 with S139/1 IgG carrying an identical Fc to CR9114 modestly increased pTyr levels in CR9114 emAb cells relative to S139/1 IgG alone (Figure 5C). These results indicate that larger, multivalent antibodies and the opsonization of virus particles with complement proteins can both increase the potency of epitope masking in some contexts, but the contribution of these factors are epitope-or antibody-dependent.

## DISCUSSION

While it is well-established that prior immunity against influenza shapes subsequent immune responses, the specific role of antibody-BCR competition is not clearly defined. Using a simplified in vitro system, we measured B cell activation and the extraction of surface-bound viral particles in the presence and absence of competing antibodies. This system recapitulates aspects of epitope masking and allows us to systematically investigate the nature of antibody-BCR competition. We find that soluble antibodies can inhibit B cell activation through both direct and indirect masking, including by binding to epitopes on other viral surface proteins. Overall, we find that membrane-proximal epitopes on HA are particularly susceptible to inhibition, presenting an additional challenge for universal vaccine design. Finally, our results show that antibody binding kinetics are a crucial determinant of epitope masking: antibodies that exchange slowly inhibit BCR activation at much lower concentrations than those that rapidly exchange even when the apparent affinities are similar. This may further disadvantage BCRs against the HA stalk, whose conserved hydrophobic residues support the binding of antibodies that can achieve notably slow dissociation kinetics (<0.001 s^-1^ for the CR9114 Fab)^23,56^.

In addition to these general trends, we also observe instances where the presence of antibodies modulates B cell activation in unexpected ways. At least one antibody we tested targeting the membrane-proximal HA anchor (FISW84) was able to potently inhibit a head-targeting BCR (C05). One possible explanation for this finding is that FISW84 forces the HA ectodomain to tilt towards the membrane, making it harder for C05 BCRs to bind to the HA apex. If correct, this model would suggest that epitope masking can also occur through allosteric mechanisms. We observe another potential example of allosteric modulation for BCRs that recognize the trimer interface epitope. While most of the antibodies we tested reduce B cell activation to variable degrees, CR9114 IgG increased phosphorylation of the FluA-20 BCR. This suggests that binding by CR9114 may destabilize the HA trimer, exposing epitopes at its interface. Interestingly, an antibody that binds to the same epitope as CR9114, 31.b.09, does not show the same effect. Understanding how antibodies influence the conformational dynamics of HA could facilitate targeting of conserved epitopes with limited accessibility.

Beyond direct competition with BCRs, opsonization of viral antigen with antibodies will likely steer the immune response through additional mechanisms. The inhibitory Fc receptor FcγRIIb has been shown to raise thresholds for B cell activation upon engagement of CD23 with viral immune complexes, leading to the production of higher affinity antibodies^57^. This effect may be dampened for B cells that recognize epitopes masked by the opsonizing antibodies. Conversely, activation of the complement cascade and subsequent opsonization of viral antigen with complement proteins can enhance B cell activation and antigen phagocytosis through the ligation of the B cell co-receptor complex^58^. While we have not examined these effects here, it is interesting to note that binding of C1 to the stalk-reactive antibody CR9114 markedly decreased phosphorylation of a competing CR9114 BCR relative to antibody alone. Future studies focused on understanding the interplay between epitope masking and complement-mediated signaling could help establish a foundation for immune complex vaccines^59^.

## ACKNOWLEDGEMENTS

This work was supported by National Institutes of Health grant R01 AI171445 and National Science Foundation CAREER Award 2238165.

The following reagents were obtained through BEI Resources: Human Convalescent Serum 001 to 2009 H1N1 Influenza A Virus, NR-18964, and Human Convalescent Serum 002 to 2009 H1N1 Influenza A Virus, NR-18965.

We thank Dr. Daved Fremont for providing the LALAPG antibody backbone, Dr. Jai Rudra for providing the Novocyte equipment, and Dr. Regina Clemens for technical insights on the calcium influx assay.

## AUTHOR CONTRIBUTIONS

Y.H. and M.D.V. designed the research; Y.H. performed the research; Y.H. and M.D.V. contributed new reagents; Y.H. and Z.G. analyzed the data; Y.H. and M.D.V. wrote the paper.

## METHODS

### Virus culture

MDCK-II cells utilized in the study for influenza virus production were obtained as authenticated cell lines (STR profiling) from ATCC. They were cultured using cell growth medium consisting of Dulbecco’s modified Eagle’s medium (DMEM; Gibco) supplemented with 10% fetal bovine serum (FBS; Gibco) and 1× antibiotic-antimycotic (Corning), and maintained under standard conditions (37°C and 5% CO_2_). Viral stocks were rescued and characterized using standard reverse genetics techniques and expanded from low multiplicity of infection (MOI) in MDCK-II cells in virus growth medium comprised of Opti-MEM (Gibco), 2.5 mg/mL bovine serum albumin (Sigma-Aldrich), 1 µg/mL L-(tosylamido-2-phenyl ethyl) chloromethyl ketone (TPCK)-treated trypsin (Thermo Scientific Pierce), and 1× antibiotic-antimycotic^60,61^. To study the effect of trypsin cleavage on FluA-20-emAb cell activation against influenza viruses, viruses were expanded from high MOI in the absence of TPCK-treated trypsin.

### B cell culture and engineering

Ramos B cells used in the study were purchased from ATCC (CRL-1596) and cultured in Iscove’s Modified Dulbecco’s medium (IMDM; Gibco) supplemented with 10% fetal bovine serum (FBS; Gibco) and 1× antibiotic-antimycotic (Corning) at 37°C and 5% CO_2_. To knock out the endogenous BCR, we cloned an sgRNA (5’-GCAGGGCACAGACGAACACG-3’) into the lentiCRISPRv2 transfer vector, and packaged VSV-G psuedotyped lentivirus in HEK293T cells. Ramos B cells (∼2.5×10^6^ cells at 0.5×10^6^ cells/ml) were transduced with concentrated lentivirus for two days. To enrich for the IgM-negative population following transduction, we pelleted cells and bound a biotinylated anti-IgM Fab fragment on ice for 10 minutes, washed in cold PBS with 0.1% BSA to remove unbound antibody, and captured IgM+ cells using streptactin magnetic beads (IBA Lifesciences). Cells that were not captured were collected and expanded. We repeated this enrichment procedure ∼2-3 times, until the percent IgM+ cells was <0.1%.

To establish B cell lines expressing single-chain BCRs, we cloned BCR sequences consisting of the light chain, a linker with three tandem strep-tags, and heavy chains with IgM or IgG1 constant regions into the pHR-SIN transfer vector. For IgM BCRs, we introduced a silent mutation in the PAM sequence targeted by our sgRNA to prevent targeting by residual Cas9 expression. Transduced B cells were subjected to sequential rounds of enrichment using streptactin magnetic beads following the manufacturer’s protocol. Sequences for BCR constructs are given in Supplementary Information.

### Protein purification and labeling

Sequences for the variable regions of antibody heavy and light chains were obtained from deposited antibody sequences and cloned into expression vectors to generate full-length human IgG1 antibodies. For both full-length IgG1 and Fab fragments, a C-terminal ybbR tag on the heavy chain is used for fluorescent labeling using SFP synthase^62^ and Fab fragments contain an additional His_6_ tag for affinity purification. For IgA antibodies, the heavy and light chains are expressed without tags and a C-terminal His_8_ tag is added to the J chain for the purification of dimeric IgA. The extracellular domain from HA is cloned by replacing the transmembrane and cytoplasmic domains with a C-terminal foldon, followed by a His_6_-tag and ybbR tag. Sequences for recombinant proteins are given in Supplementary Information.

Antibodies and HA are expressed in HEK-293T cells for 6-7d following transfection at >70% confluency. Cells are cultured in Opti-MEM, antibiotic-antimycotic, and 2% FBS (for HA, IgA, and Fab fragments) or without FBS (for full-length IgG1). His-tagged proteins are purified from cell culture supernatants using Ni-NTA agarose beads (Thermo Scientific Pierce) and IgG1 antibodies are purified using protein A agarose beads (Thermo Scientific Pierce). Human convalescent sera were provided by BEI Resources (NR-18964 and NR-18965 for Serum 001 and Serum 002, respectively) and purified with protein A agarose beads (Thermo Scientific Pierce) prior to use in antigen extraction assays. Anti-IgG Fabs were prepared by AffiniPure Fab Fragment Goat Anti-Human IgG (Jackson ImmunoResearch Laboratories) and fluorescently labeled with NHS dyes.

### Virus immobilization

Glass-bottom 96-well plates (Cellvis) were prepared for antigen extraction experiments by coating with 0.18 mg/mL biotinylated BSA in PBS for 2 hours at room temperature. The imaging plate was then washed with PBS twice and incubated with streptavidin (Invitrogen) at 25 ug/ml in PBS for 2 hours at room temperature. Next, the imaging plate was washed with PBS twice and incubated with 25 µg/mL biotinylated *Erythrina cristagalli* lectin (ECL; Vector Laboratories) at room temperature for 2 hours. Finally, the imaging plate was washed with PBS up to 5 times and stored at 4°C until use.

### Antigen extraction assay

Viruses freshly expanded from MDCK cells were immobilized onto ECL-treated plates by centrifugation at 1500×g for 10 min. Unbound viruses were removed by washing with warm cell culture medium up to ten times. Fluorescent C05, FISW84, or FI6v3 Fab were diluted to 1nM for visualization as needed. Virus samples were imaged on a Nikon Ti2 microscope equipped with a CSU-X1 spinning disk and Tokai Hit stage-top incubator using a 40×, 1.3 NA oil objective. At least five fields of view per well were imaged before adding the emAb cells and incubating at 37°C for 1h. Prior to experiments with virus particles, emAb cells were treated with 0.1 U/mL sialidase from *Clostridium perfringens* at 37°C for 30 mins to minimize interactions between viral HA and sialic acid on the B cell surface. The wells with virus and B cells were then imaged again to determine the fraction of virus particles that were removed from the coverslip. Percentage virus reduction is calculated by dividing the changes in particle number by total virus number before adding B cells.

### Immunofluorescence

Viruses were immobilized onto ECL-treated plates, incubated with fluorescent non-competing CR9114, FISW84, or C05 Fab with or without competing antibodies at 2x the desired final concentration in a volume of 100ul. After 30 minutes at 37°C, 3x10^5^ emAb cells in 100ul cell culture medium containing 0.2 U/mL *Cp*NA (Sigma-Aldrich) were added to plates with virus-antibody complexes for 30 minutes at 37°C. To probe for phosphotyrosine, we gently wash with PBS and incubate at room temperature for 15 min in a 1% PFA/PBS solution. After washing twice with PBS, we permeabilize with 0.1% Tween-20 for 15 min at room temperature before washing twice with PBS. After blocking with 10mg/ml BSA in PBS for 30 min at room temperature, we add anti-pTyr antibodies (P-Tyr-1000 MultiMab Rabbit mAb mix, Cell Signaling Technology) at 1:400 dilution and incubate overnight at 4°C or at room temperature for 2h. After washing with PBS, cells are incubated with Goat-anti-Rabbit IgG conjugated with AlexaFluor 647 (ThermoFisher Scientific, A-21244) at 4 µg/mL and incubated at room temperature in the dark for 1h before imaging with a 20x, 0.75-NA or 40x, 1.3-NA objective.

### Calcium influx assay

Influenza viruses (A/California/04/2009) were immobilized onto ECL plates, visualized via fluorescently labelled FISW84 Fab, and incubated with or without 60nM CR9114 IgG antibodies. CR9114-emAb cells were incubated with calcium sensitive dye from a Fluo-4 Calcium Imaging Kit (Invitrogen) at 1:1000 dilution for 30 min at 37°C. Cells were washed with warm cell culture medium twice to remove extra dye and kept on ice until experiment. The cells were warmed up at 37°C before adding to the well and imaging immediately at 5s per frame for 10 min using a 60x, 1.40-NA oil objective. At least 10 cells that encountered virus particles on the glass-bottom plate were randomly selected to extract its median intensity over time for each experimental condition. Using a custom Python script, baseline fluorescence signals from the first frame were subtracted from the rest of the trajectory, and each trajectory was registered to the first frame where the median intensity reached 10% its maximum value.

### Antibody competition assay

Influenza virus particles were immobilized on ECL-treated plates and incubated with HA-or NA-specific antibodies for 30 min at room temperature before adding a second antibody while maintaining the same concentration of the initial antibody. Images of virus particles and bound antibodies were collected using a 60x, 1.40-NA objective and analyzed using segmentation in Nikon Elements Software.

### Supported lipid bilayers

Glass coverslips were cleaned with piranha (3:2 mixture of sulfuric acid and 30% hydrogen peroxide) and coated with a 1 mg/ml suspension of small unilamellar vesicles (SUVs) consisting of 95.8 mole-percent DOPC (Avanti Research, #850375), 0.2% Atto 390 DOPE (ATTO-TEC), 2% 18:1 PEG2000 PE (Avanti Research, #880130), and 2% 18:1 DGS-NTA(Ni) (Avanti Research, #790404) overnight at RT. The wells were washed with PBS ∼10 times, incubated with fluorescently labelled HA ectodomain (A/Hong Kong/1968) at room temperature, and washed again with PBS at least 10 times. For experiments with competing antibodies, the wells were incubated with soluble antibodies at 37°C for 1h prior to adding emAb cells. Images of emAb cells on supported lipid bilayers were taken at 30 min using a 60×, 1.40 NA oil objective.

### Measuring cell-surface BCR expression

Engineered B cells (10^6^ cells) were washed twice with PBS and resuspended in 100 µL of chilled PBS with 10nM fluorescent anti-IgG Fab. After labeling for 10 min on ice, emAb cells were analyzed using a NovoCyte (ACEA Biosciences, Inc.). Data was collected for at least 0.5x10^6^ cells per sample and gated for analysis using uniform thresholds across samples. Fluorescence intensity of single B cells was analyzed using custom Python analysis script to obtain the median intensity values.

### Statistics and Replicates

Statistical analysis was performed using Python. No statistical methods were applied to predetermine sample size. Statistical tests used are indicated in each respective figure legend. Biological replicates are defined as cells separately infected/transfected/treated and assayed in separate wells as indicated.

## Data Availability

The manuscript presents analyzed data with some raw images for demonstration purposes. Raw images will be uploaded to Image Data Resource (https://idr.openmicroscopy.org/) upon publication. Analysis code and code used for figure generation will be uploaded to GitHub.

---

**Figure S1.**
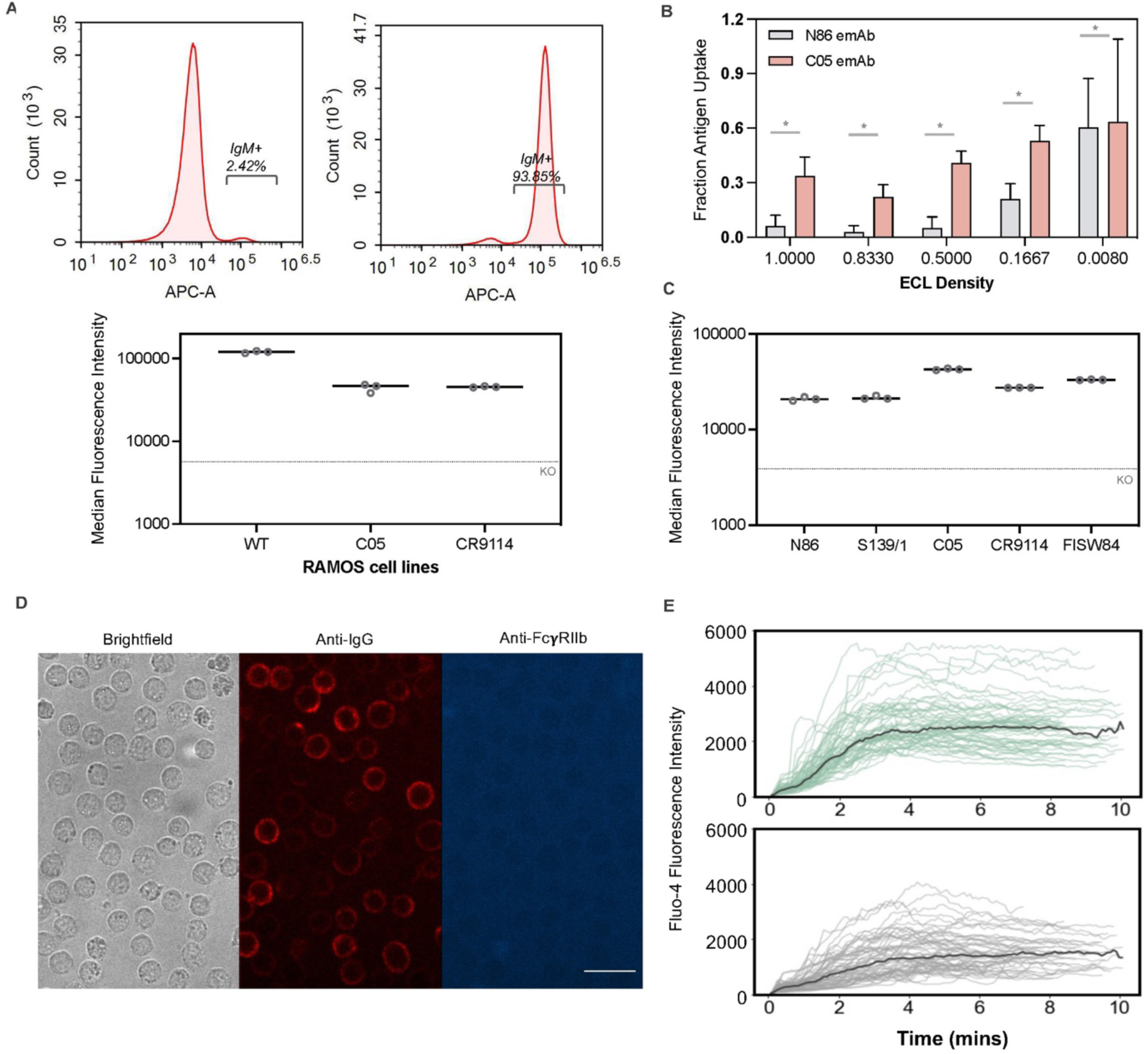
Engineering B cells for imaging-based assays of BCR engagement and activation. (A) Top: Representative results from flow cytometry using fluorescently labelled anti-IgM Fab for knock-out Ramos cells (top) and wildtype Ramos cells. “IgM+” represents the gating method for IgM-positive B cells and is standardized across conditions. Bottom: Median fluorescence intensities of cell surface BCRs (IgG isotype) on N86-emAb, S139/1-emAb, C05-emAb, CR9114-emAb, and FISW84-emAb cells. (B) Comparison of antigen extraction by non-specific N86-emAb cells and influenza-specific C05-emAb cells at varying fractional densities of surface ECL. Data are combined from three biological replicates containing five fields of view each. *P*-values are determined by independent t test. (C) Median fluorescence intensity of cell surface BCRs (IgM isotype) on wildtype, C05-emAb, and CR9114-emAb cells. Data are combined from three biological replicates containing >0.5×10^6^ cells each. (D) Representative images of CR9114-emAb cells labeled with anti-IgG Fab and a monoclonal antibody against FcγRIIb (S18005H). (E) Quantification of calcium-sensitive Fluo-4 intensity for CR9114-emAb cells presented with A/California/04/2009 virus particles in the presence or absence of competing CR9114 IgG. Fluorescence trajectories are aligned to the first frame at which they reach 10% of their final intensity value. Bold curves indicate median intensity values at each time point. Data are combined from three biological replicates containing at least 10randomly sampled emAb B cells per replicate.

**Figure S2.**
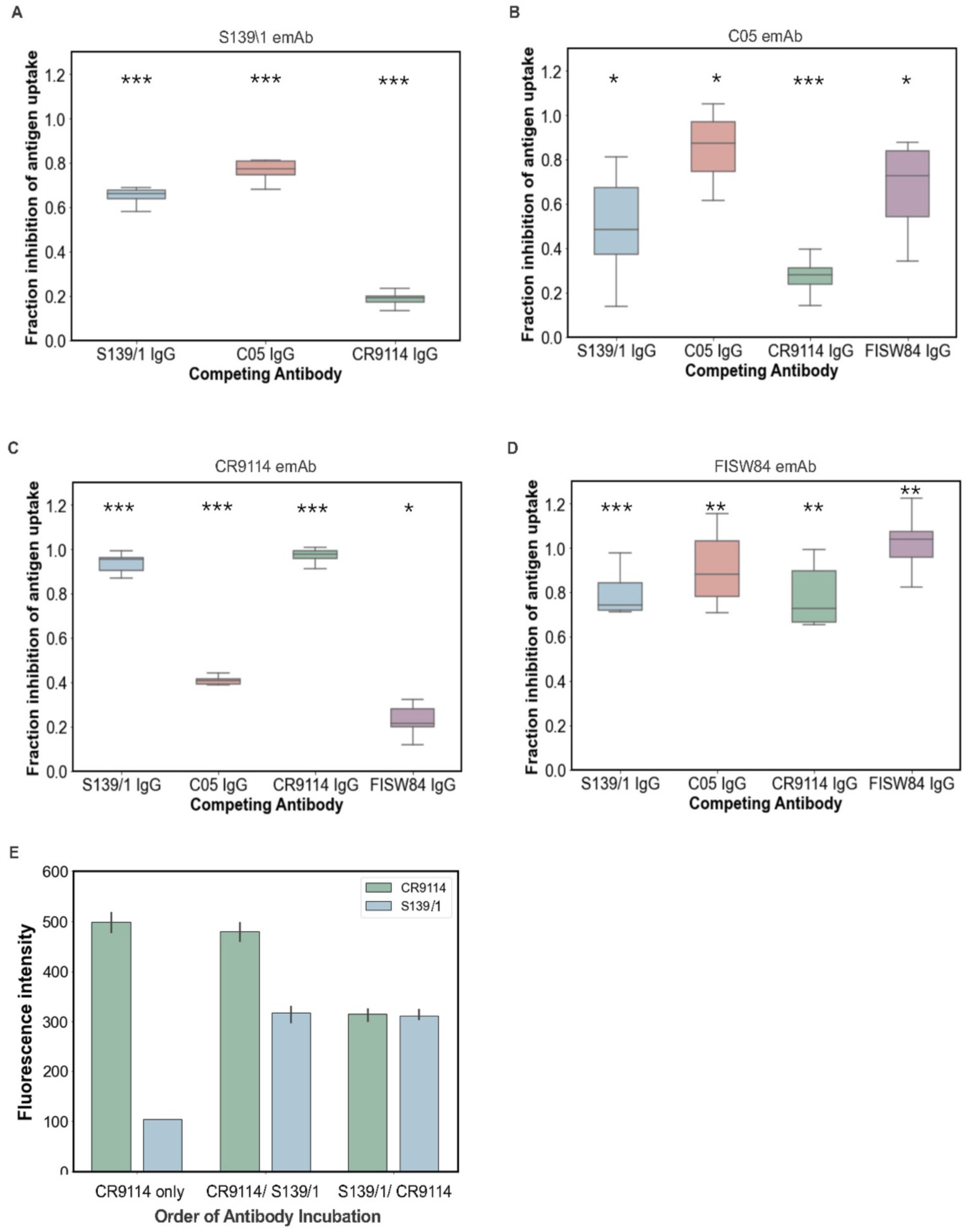
B cell antigen uptake is modulated by antibodies targeting directly and non-directly competing epitopes. (A-D) Fraction inhibition of antibody uptake by directly and non-directly competing antibodies at 60nM against emAb cells. Values of ∼1 indicate complete inhibition, whereas values of ∼0 indicate no inhibition. Data are combined from three biological replicates containing five fields of view each, and are plotted in matrix form in Figure 3C. *P*-values are determined by independent t test against the normalized antibody-free condition (=0, not shown in the figure). (E) Quantification of antibody binding to A/WSN/1933 viruses in different sequences: CR9114 IgG only; CR9114 before S139/1; or CR9114 after S139/1. Both antibodies are tested at 60nM.

**Figure S3.**
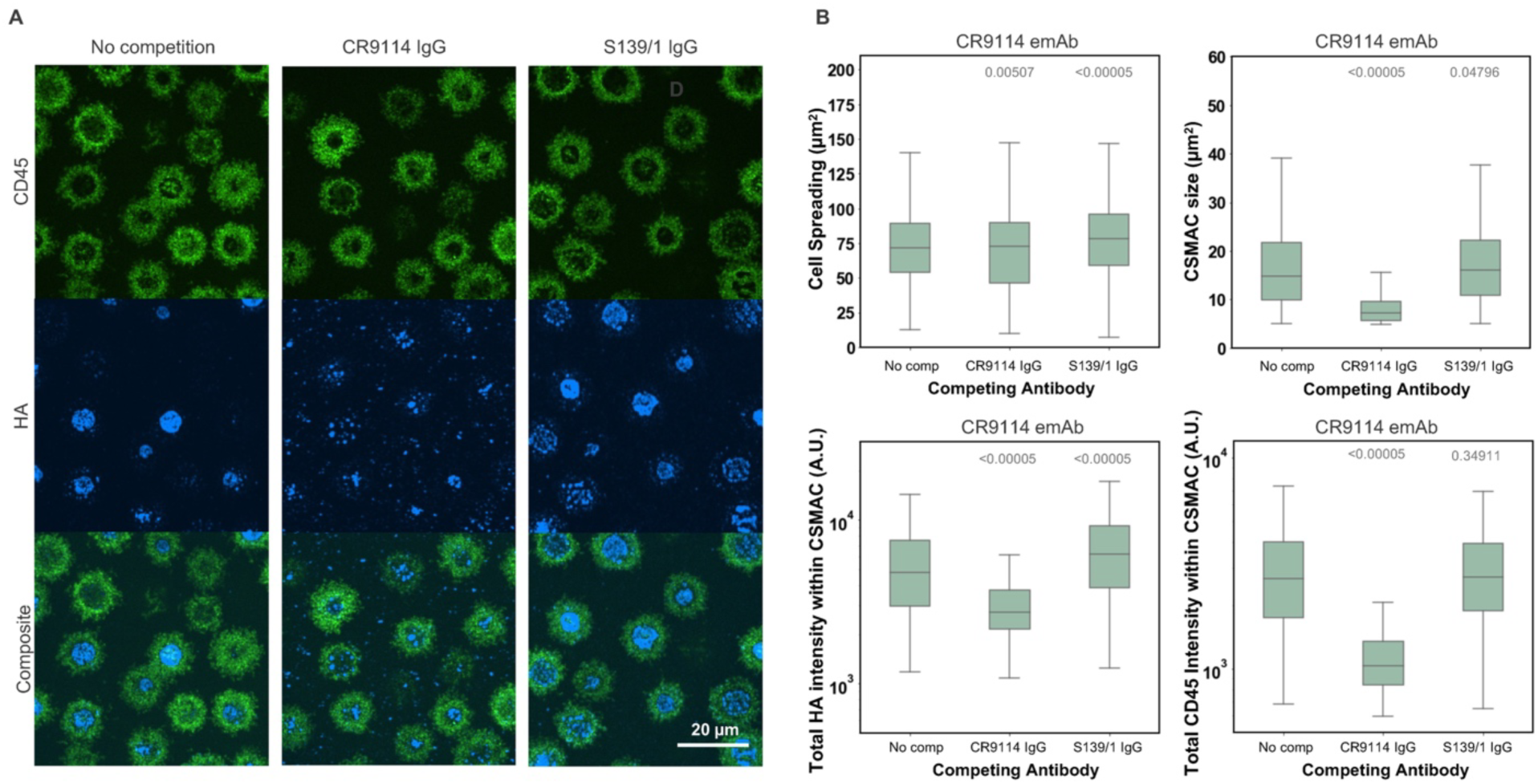
Epitope masking on fluid lipid bilayers. (A) Representative images of CR9114-emAb cells accumulating HA (A/Hong Kong/1968) presented on a supported lipid bilayer in the presence or absence of competing CR9114 or S139/1 IgG, both at 60nM. (B) Quantification of CR9114-emAb cell synapses in the presence or absence of competing CR9114 or S139/1 IgG. Plots show cell spreading and cSMAC size (top row) and HA accumulation and CD45 signal (bottom row). Data points represent measurements for individual B cells segmented either by the entire cell periphery or the cSMAC region. Data are combined from two biological replicates containing 10 fields of view each. *P*-values are determined by Kolmogorov-Smirnov test. Statistical comparisons in panel B are to the condition without competing IgG.

**Figure S4.**
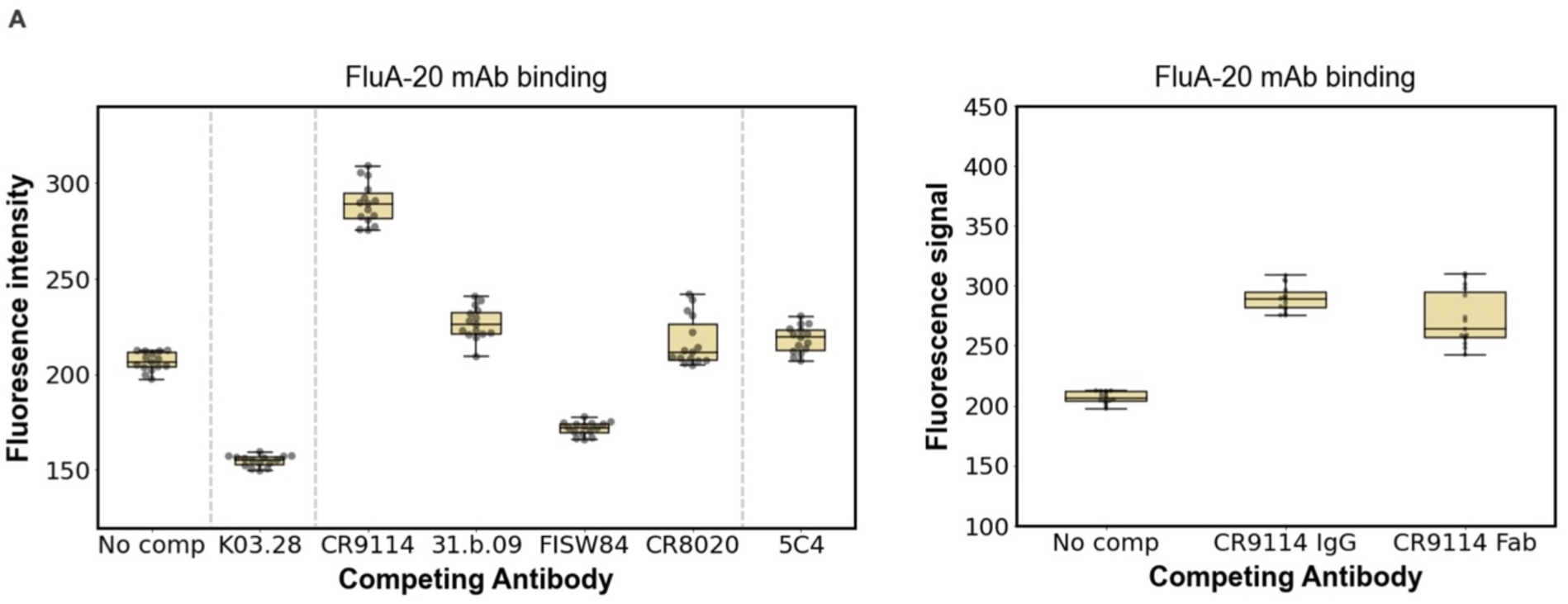
Hemagglutinin stability varies upon binding of soluble antibodies. (A) Quantification of FluA-20 IgG binding to A/California/04/2009 viruses in competition with other IgG (at 60nM) and Fab (at 120nM). Data are combined from three technical replicates containing five fields of view each. *P*-values are determined by independent t test using media values from each technical replicate.

## Supplementary Movie Captions

**Movie S1: Influenza A virus extraction from coverslips by CR9114-emAb cells.** Timelapse microscopy of B cells expressing a BCR (shown in red) derived from CR9114 engaging with A/WSN/1933 virus particles (shown in blue). Images are collected at 30s intervals for 30 minutes.

**Movie S2: Influenza A virus extraction from coverslips by CR9114-emAb cells is blocked by competing IgG.** Timelapse microscopy of B cells expressing a BCR (shown in red) derived from CR9114 engaging with A/WSN/1933 virus particles (shown in blue) pre-incubated with 10 nM CR9114 IgG. Images are collected at 30s intervals for 20 minutes.

**Movie S3: Synapse formation for CR9114-emAb cells presented with HA on a supported lipid bilayer.** Timelapse microscopy of CR9114 emAb B cells presented with His-tagged HA (from A/Hong Kong/1968; shown in magenta) on a supported lipid bilayer. CD45 is shown in green and the BCR is shown in blue. Images are acquired at 60s intervals for 43 minutes. Scale bar = 10µm.

